# Desmoglein-3 modulates p38MAPK and ERK signaling responses through the mechano-sensitive channel Piezo1

**DOI:** 10.64898/2026.05.11.723746

**Authors:** Karen Leal-Fischer, Henriette Franz, Katarzyna Buczak, Aude Zimmermann, Volker Spindler

## Abstract

**Background:** Skin is constantly exposed to mechanical forces such as pressure and friction, which need to be sensed and buffered to ensure tissue homeostasis and barrier function. Desmosomes are essential for epidermal integrity, but their role in converting mechanical cues into cellular signaling responses are not well understood.

**Methods:** Here, we combine proteomics and shear-stress assays with live-cell reporters to investigate how desmosomes modulate stress-kinase pathways in keratinocytes.

**Results:** We show that the desmosomal adhesion molecule DSG3 is essential not only for cell-cell adhesion but also for modulating p38MAPK and ERK signaling. Loss of DSG3 disrupts mechanotransduction-related protein networks, including the expression of the mechanosensitive channel Piezo1. Under static conditions, DSG3 dampens ERK activity via Piezo1-dependent mechanisms, whereas DSG3 suppresses p38MAPK activity through an independent mechanism. In contrast, DSG3 is required to trigger an activation of both ERK and p38MAPK in response to shear stress in a Piezo1-dependent manner. Experiments with domain-specific DSG3 mutants demonstrate that cell cohesion and signaling responses are partially uncoupled, while maintaining DSG3 tail integrity was crucial for p38MAPK and ERK responses.

**Conclusion:** These findings demonstrate that DSG3 independently coordinates adhesion and mechanotransduction in a domain-specific manner, providing novel insights into how DSG3 contributes to epithelial integrity under dynamic mechanical environments.

## Background

Multicellular organisms rely on cell-cell adhesion to form resilient tissues such as skin, which serve as essential barriers against external threats and maintain body homeostasis (1, 2). Epidermis, the outermost layer of skin, is continuously exposed to mechanical stresses, including pressure, friction and tension (3–5). To withstand these stresses, epidermis relies on the function of desmosomes which ensure strong cell adhesion in tissues subjected to high mechanical load (6, 7). They consist of transmembrane cadherin-type adhesion proteins, desmogleins (DSG) and desmocollins (DSC), the intracellular adapters plakoglobin (PG), plakophilins (PKPs), and desmoplakin (DSP). The latter links the complex to the intermediate filament cytoskeleton (8, 9). Desmoglein intracellular domains are composed of an intracellular anchor (IA), an intracellular cadherin-like segment (ICS), a linker domain (LD), a repeat unit domain (RUD), and a desmoglein terminal domain (DTD). The ICS domain is known to bind PKPs and PG, while the RUD domain has been shown to stabilize DSG2 at the membrane (10). Nevertheless, the functions of the intracellular tail with regard to cellular signaling events remain only partially understood. Disruption of desmosomal adhesion leads to severe diseases mainly affecting skin and heart (9, 11). Knockout studies and observations from disease models demonstrated that specific desmosomal cadherins such as desmoglein 3 (DSG3) and desmoglein 1 (DSG1) modulate the activity of MAPK pathways, in particular these of p38MAPK and ERK as well as cellular calcium flux (12–17). Moreover, desmosomal adhesion was shown to strengthen under mechanical load, thereby increasing tissue resilience to forces (4, 18, 19). A recent study using FRET-based sensors demonstrated that DSG3 is under tension at the membrane, which is lost under desmosome disease conditions by a mechanism involving RhoA activation (20). These findings exemplify that, beyond their structural role, desmosomes integrate cell adhesion with signaling pathways (21, 22). Interestingly, while adherens junctions are established mechanotransducers (23–25), it is only partially understood whether desmosomes can sense force and how they specifically trigger signaling responses. For example, while it was shown that DSG3 can interact with active p38MAPK in disease settings (26–29), the specific mechanisms by which loss of desmosome function leads to p38MAPK activation are still unresolved.

To address these aspects, we use kinase translocation reporters to monitor real-time kinase activity in living keratinocytes under static and dynamic conditions to investigate the dependency of signaling responses on DSG3 and its specific intracellular domains. We show that DSG3 modulates the activity of the mechanosensitive channel Piezo1, which is important for ERK and p38MAPK responses to shear stress. These results demonstrate a role for DSG3 in mechanosensitive MAPK signaling and further add mechanistic insights to the understanding of desmosomal cadherins integrating cell-cell adhesion and cellular responses.

## Results

### DSG3 intracellular domains are essential to dampen p38MAPK activity but dispensable for ERK suppression in static conditions

To investigate the role of desmoglein-3 (DSG3) in keratinocyte mechanotransduction, a knockout model (DSG3-KO) was generated in the human HaCaT keratinocyte cell line using the CRISPR/Cas9 system. Cells transduced with a non-targeting guide RNA served as controls (CTRL). We first examined the effects of DSG3 loss under static conditions. Deletion of DSG3 resulted in a marked reduction of desmoplakin (DSP) at the cell membrane and reduced levels of plakoglobin and DSG2 (Fig.1A, B, Fig. S1A, B), indicating fewer or smaller desmosomes. DSG3-KO cells exhibited reduced cortical keratin localization and impaired F-actin distribution (Fig. 1C, D, Fig. S1C), and displayed a significantly increased number of fragments after applying shear stress to a detached monolayer as an indication of impaired cell-cell adhesion (Fig. 1E). Loss of DSG3 led to increased phosphorylation of p38MAPK and ERK (p-p38MAPK, pERK) (Fig. 1F, G) consistent with previous studies (28).

**Figure 1:**
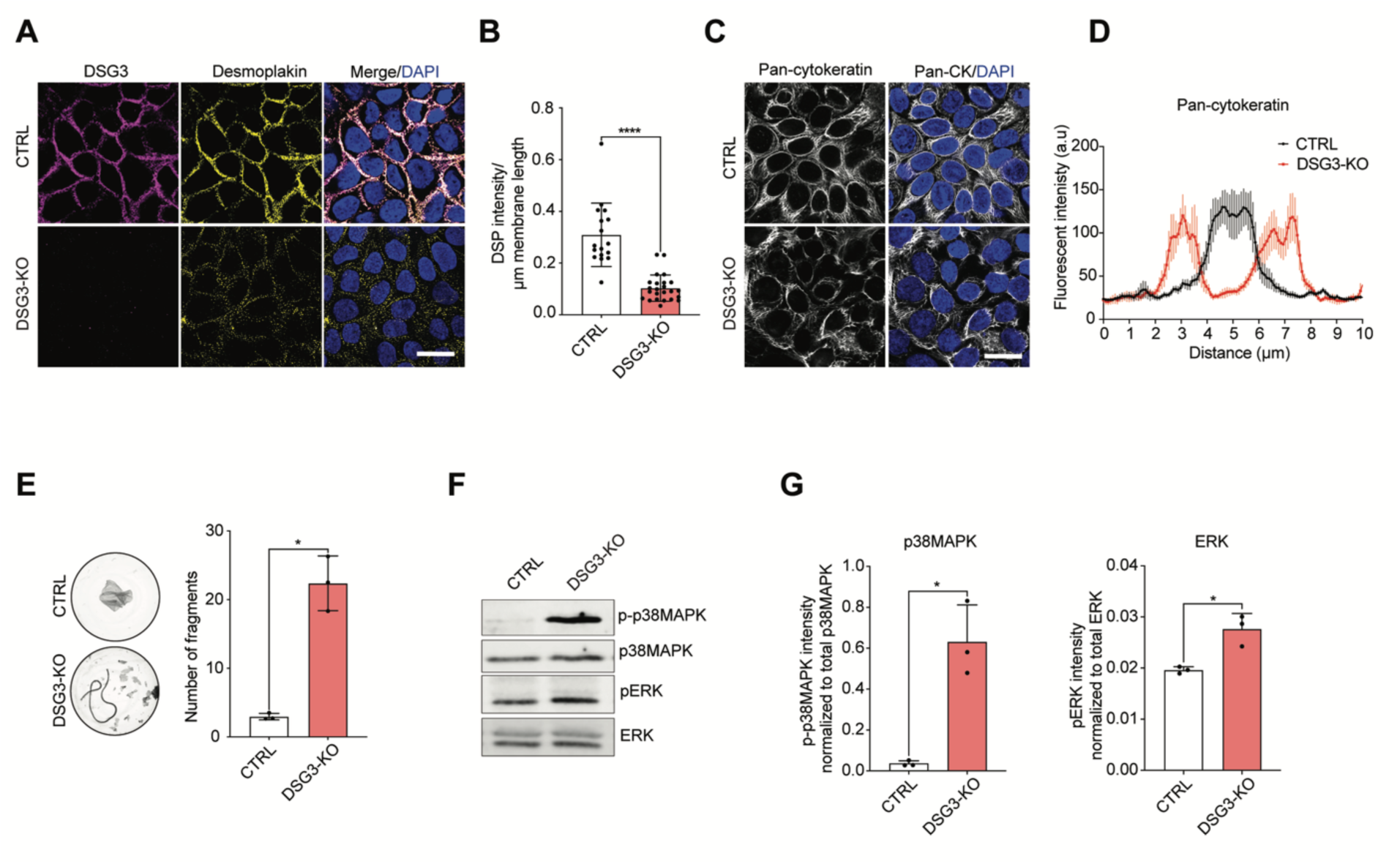
DSG3 modulates cell adhesion and MAPK signaling. **A**. Immunofluorescence staining for DSG3 and desmoplakin (DSP) of CTRL and DSG3-KO HaCaT keratinocytes. Scale bar 10 μm. **B**. Quantification of DSP intensity over the corresponding membrane length (μm), Mann-Whitney test. Each data point represents one cell from three independent experiments in total. **C**. Immunofluorescence staining for Pan-cytokeratin (pan-CK) of CTRL and DSG3-KO HaCaT keratinocytes. Scale bar 10 μm. **D**. Analysis of pan-CK staining by quantifying the intensity (a.u) perpendicular to cell borders over a distance of 10 µm. **E**. Dispase-based dissociation assay of CTRL and DSG3-KO cells. Representative images and quantifications of N = 3 are shown, Welch’s t-test. **F-G**. Western blot and representative quantification of phosphorylated p38MAPK (p-p38MAPK) and ERK (pERK) in CTRL and DSG3-KO HaCaT keratinocytes. Each dot represents biological replicates. Welch’s t-test.

**Figure S1:**
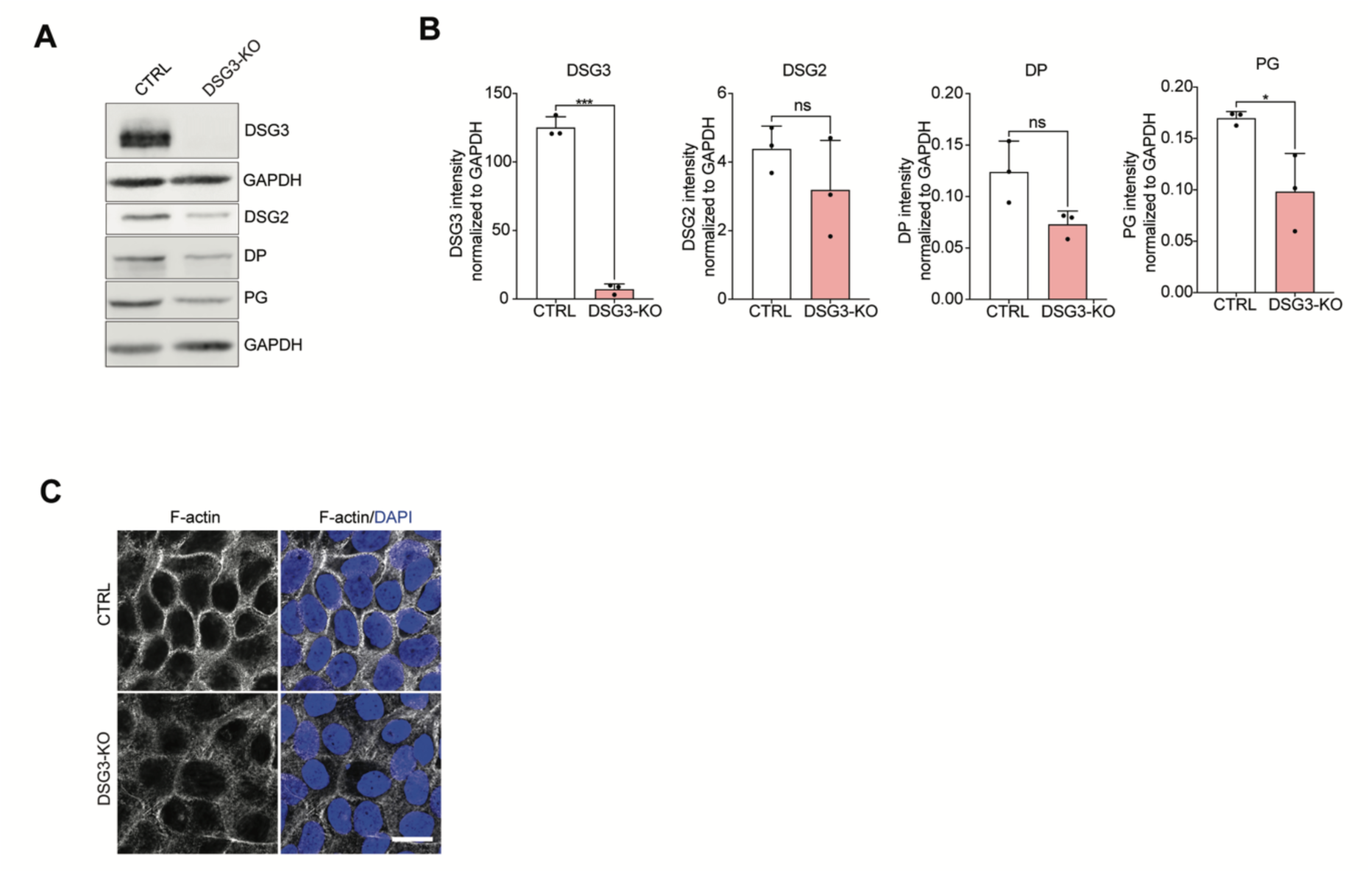
DSG3-KO characterization. **A-B**. Western blot and representative quantification of DSG3, DSG2, DP, and PG in CTRL and DSG3-KO HaCaT keratinocytes. Each dot represents biological replicates. Unpaired t-test. **C**. Immunostaining for F-actin in CTRL and DSG3-KO HaCaT keratinocytes. Scale bar 10 μm. Images are representative of three biological replicates.

As it was previously shown that p38MAPK can either directly or indirectly interact with DSG3 (26, 28, 30–33), we tested the hypothesis that the intracellular tail domains of DSG3 are important to modulate p38MAPK and ERK responses. Full-length DSG3 and mutants lacking specific domains (IA, ICS, LD, RUD) were generated and transduced into DSG3-KO cells (Fig. 2A). All mutants except DSG3-ΔLD-RFP (Fig. S2A) localized to the membrane. To study real-time kinase activity, fluorescent kinase translocation reporters (KTRs) for ERK and p38MAPK (34, 35) were transduced and evaluated by live-cell imaging (Fig. 2B). These reporters localize to the nucleus or the cytosol, depending on the activity of the respective kinase. A semiautomated image analysis pipeline was established to determine the cytosolic-to-nuclear ratio of the KTR fluorescent signal (Fig. S2B). DSG3-KO cells exhibited elevated and, within the imaging intervals tested, stable p38MAPK and ERK activity (Fig. 2C-F) under static conditions, consistent with the increased phosphorylation levels observed by western blot (WB). Re-expression of full-length DSG3 significantly reduced basal p38MAPK activity and fully restored pERK levels (Fig. 2C-F). Basal p38MAPK activity remained elevated in all mutants, suggesting that all intracellular domains are required to dampen p38MAPK activity. Interestingly, basal ERK activity was restored by all DSG3 mutants, including DSG3-ΔLD, which lacked membrane localization.

**Figure 2:**
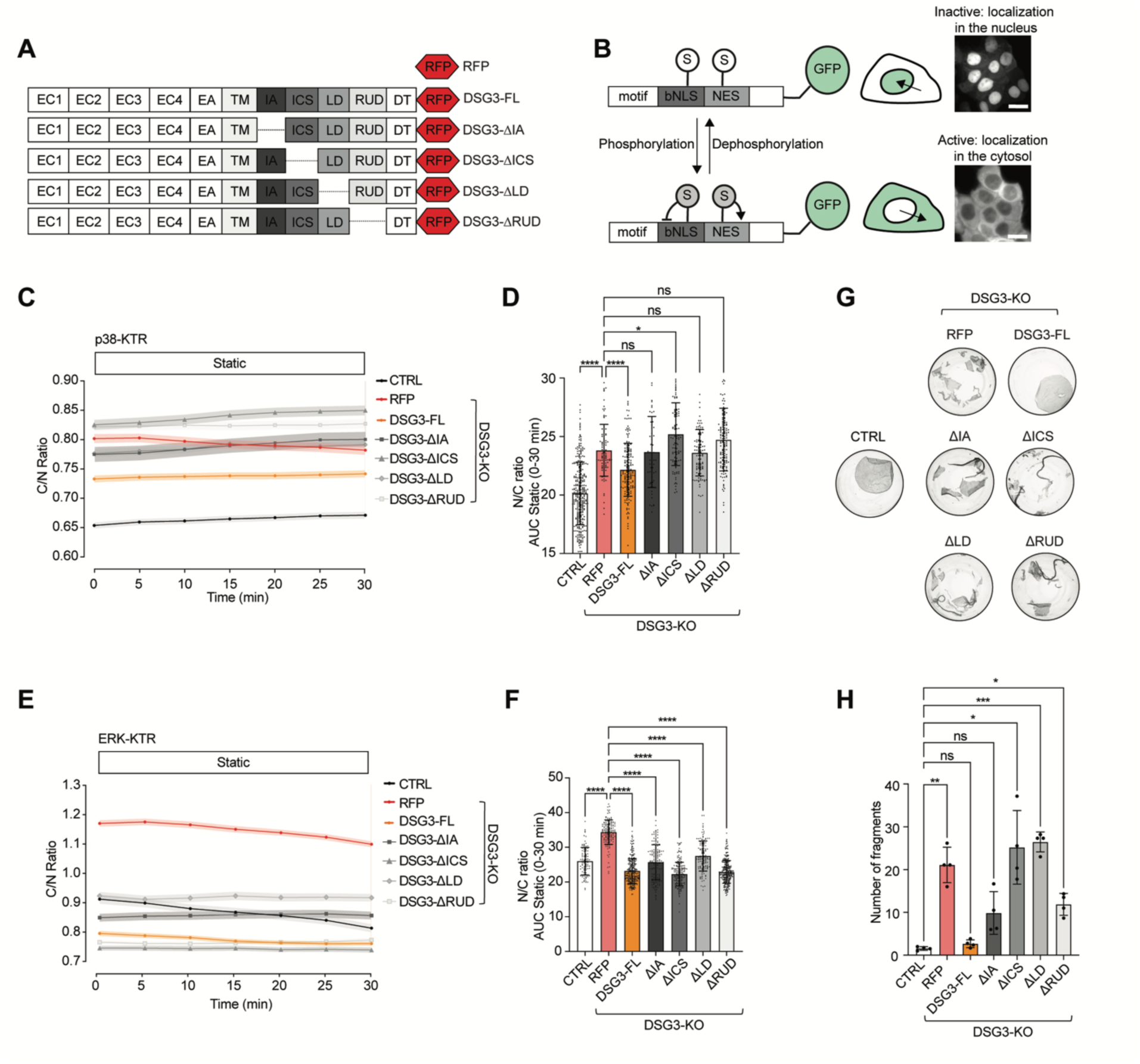
DSG3 intracellular domains are essential for basal p38MAPK activity. **A**. Schematic of RFP and RFP-tagged DSG3 constructs: full-length desmoglein 3 (DSG3-FL) and domain-specific mutants where the intracellular anchor (DSG3-ΔIA), intracellular cadherin-like segment (DSG3-ΔICS), linker domain (DSG3-ΔLD), and the repeat unit (DSG3-ΔRUD) domains were deleted. **B.** Schematic representation of Kinase Translocation Reporters (KTRs): a substrate recognition motif fused to a bipartite nuclear localization signal (bNLS), a nuclear export signal (NES), and a Green fluorescent protein (GFP). The sensor translocates from the nucleus to the cytosol upon kinase phosphorylation. Modified from Kudo et al., Nature Protocols, 2018. **C, D**. p38MAPK and ERK activity of CTRL, DSG3-KO, and DSG3 mutants expressing the KTR sensors depicted as the C/N ratio (cytoplasmic over nuclear intensities) over time under static conditions. Data are represented as the mean ± SEM from 70-295 cells of three independent experiments. **D, F**. Quantification of the area under the curves (AUC) over 30 min shown in C and E. N=3, One-way ANOVA Dunn’s multiple comparison test. **H**. Dispase-based dissociation assay of CTRL, DSG3-KO RFP, DSG3-FL, and DSG3 mutants. Representative images of cell monolayers after shearing are shown. **I**. Quantification of the number of fragments generated, N=4, Dunn’s multiple comparison test.

To correlate MAPK signaling and cell cohesion, we also evaluated the effect of the mutants in dispase-based dissociation assays (Fig. 2G, H). Re-expression of full-length DSG3-RFP in DSG3-KO cells restored cell cohesion by preventing monolayer sheet fragmentation (Fig. 2 H, I). Expression of DSG3-ΔIA and DSG3-ΔRUD mutants partially prevented fragmentation, whereas DSG3-ΔICS and DSG3-ΔLD mutants failed to recover adhesion, as fragmentation levels were comparable to those in DSG3-KO cells. These results suggest that at least the ICS domain and proper membrane localization of DSG3 are essential for maintaining cell cohesion.

Together, these results demonstrate that DSG3 suppresses both p38MAPK and ERK signaling under static conditions. However, this suppression did not correlate with global cell cohesion and required intracellular DSG3 domains only in the case of p38MAPK. In contrast, ERK suppression was independent of intracellular DSG3 domains. This suggests that DSG3 regulates p38MAPK and ERK pathways by differential mechanisms.

**Figure S2:**
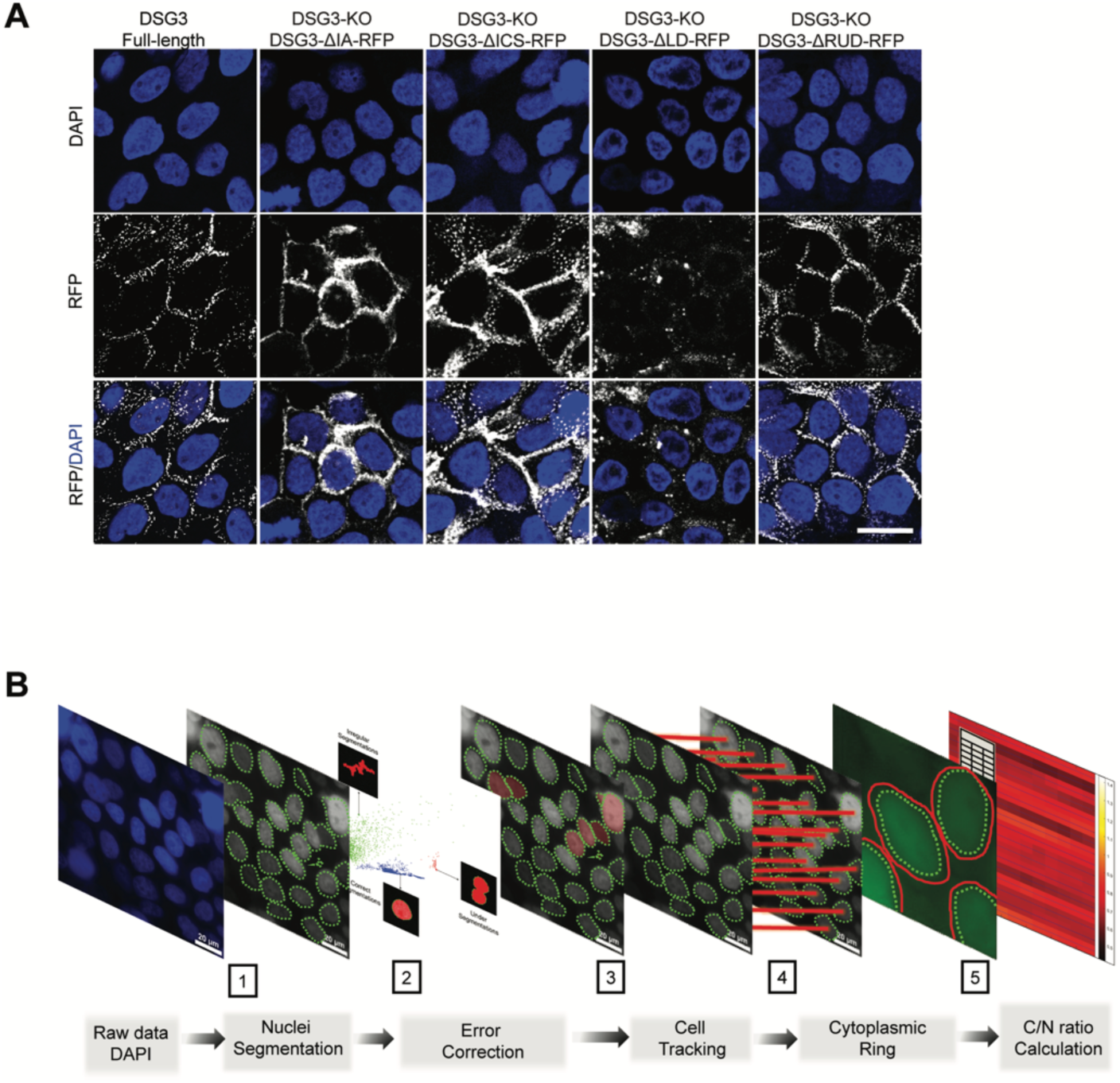
DSG3 mutants and automated live-cell KTR data analysis. **A**. Immunostainings of DSG3-KO HaCaT keratinocytes expressing RFP-tagged DSG3-FL, DSG3-ΔIA, DSG3-ΔICS, DSG3-ΔLD and DSG3-ΔRUD. Scale bar 10 μm. **B**. Workflow scheme of the key steps of MATLAB-based eDETECT automated time-lapse data analysis of the KTR sensors: **1**. Raw Hoechst staining images are used to perform nuclei segmentation. **2**. eDetect provides a fast segmentation error detection and correction of a group of errors with batch operations. **3**. Tracking of single-cell movement and position from long-term time-lapse imaging datasets. **4.** A 5-pixel-wide peri-nuclear cytoplasmic ring is used to determine the cytoplasmic regions of each cell. **5**. Measurements of the mean fluorescence intensities of the KTR construct in the nucleus and the cytoplasm, and calculation of the relative cytoplasmic versus nuclear intensity ratio (the C/N ratio), which is used as a proxy for the kinase activity in living single cells.

### Changes in the proteome in response to loss of DSG3

To better understand the different kinase activity profiles in CTRL and DSG3-KO cells, we performed mass spectrometry-based proteomic analysis to identify differentially expressed proteins in the DSG3-KO cells. Global proteomics revealed extensive remodeling of proteins involved in cell-cell junctions, cytoskeletal organization, and signal transduction (Fig. 3A-C). In addition to DSG3, several other desmosomal and junctional components, including DSG2, DSC3, plakoglobin (JUP), plakophilins (PKP2 and PKP3), and tight junction-associated proteins (TJP1 and TJP3), were significantly downregulated (Fig. 3A). In addition, mechanotransduction-related proteins such as PIEZO1, YAP1, PLEC, ICAM1, and ITGB6 were significantly altered, suggesting that loss of DSG3 may impact the ability of the keratinocytes to sense and respond to mechanical cues. STRING (Search Tool for Retrieval of Interacting Genes/proteins)-based pathway analysis of differentially expressed proteins showed significant upregulation of pathways related to cellular responses to stress and stimuli (Fig. 3B). Key components of the LINC complex (Linker of Nucleoskeleton and Cytoskeleton) including Sun1, integrins, emerin, plectins and myosin-10 were also downregulated in DSG3-KO cells (Fig.3C). The LINC complex is known to connect the cytoskeleton to the nuclear envelope, thus mediating force transmission from the extracellular matrix to the nucleus (36, 37). In addition, multiple signaling pathways were affected, including MAPK-associated pathways, as shown by upregulated expression of MAP2K2, STK24, MARK2, PAK2, and ROCK1. Together, these data indicate that DSG3 loss not only compromises desmosomal integrity but also triggers generalized changes in mechanotransduction pathways, likely contributing to altered cellular mechanics, adhesion strength, and stress responses.

**Figure 3:**
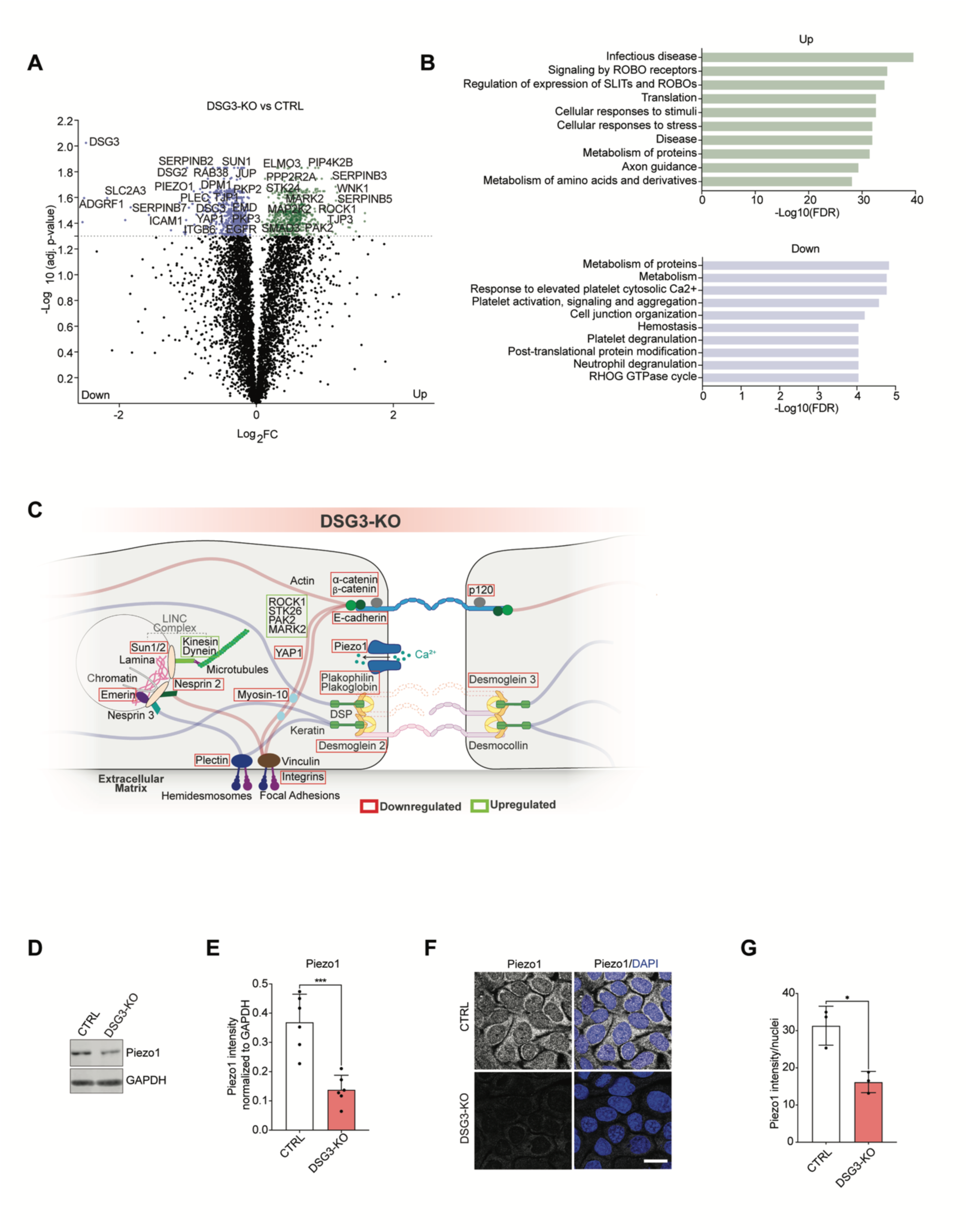
Proteomic analysis revealed Piezo1 as a downregulated mechanotransducer in DSG3-KO cells. **A.** Volcano plot showing differential protein expression in DSG3-KO HaCaTs compared to CTRL. -Log10 adj. p-values above 1.3 (0.05) were considered significant (gray dotted line). Purple dots indicate proteins that were significantly downregulated in DSG3-KO cells, whereas green dots indicate proteins that were significantly upregulated. **B.** STRING-based biological pathway analysis of significantly modulated proteins (Purple: downregulated pathways, green: upregulated pathways). The X-axis represents −log10 of the FDR values, with a cutoff of 0.05 for significance. **C.** Schematic of representative up (green) and downregulated (red) proteins in the DSG3-KO cells. **D**. Western blot images of Piezo1 and GAPDH expression in CTRL and DSG3-KO cells. **E.** Quantification of Piezo1 intensity normalized to GAPDH intensity, N=6, Welch’s t test. **F.** Immunofluorescence staining for Piezo1 in CTRL and DSG3-KO cells. Scale bar 10 μm. **G**. Quantification of Piezo1 intensity over the number of nuclei, N=3, Welch’s t test.

Piezo1 is known to trigger calcium influx in response to mechanical stress (38). Due to its key role in mechanotransduction, we focused on Piezo1. The reduced Piezo1 expression in the DSG3-KO, detected by proteomics, was confirmed by WB (Fig. 3D, E) and immunostaining (Fig. 3F, G).

### Piezo1 activity selectively regulates ERK signaling but not p38MAPK in static conditions

To investigate the relationship between PIEZO1 and p38MAPK/ERK signaling, calcium imaging was performed under static conditions. Interestingly, DSG3-KO cells exhibited significantly reduced steady-state intracellular calcium levels compared with control cells (Fig.S3A, B). Live-cell imaging using the KTR sensors in DSG3-KO cells revealed that pharmacological activation of Piezo1 with Yoda1 did not significantly affect p38MAPK activity (Fig.S3C-E), whereas ERK activity was significantly decreased compared to DMSO-treated controls (Fig.S3F-H). These findings indicate that Piezo1 activation selectively suppresses ERK signaling without modulating p38MAPK activity. Conversely, inhibition of Piezo1 using GsMTx-4 showed a trend toward increased ERK activity (Fig.S3I-K).

**Figure S3:**
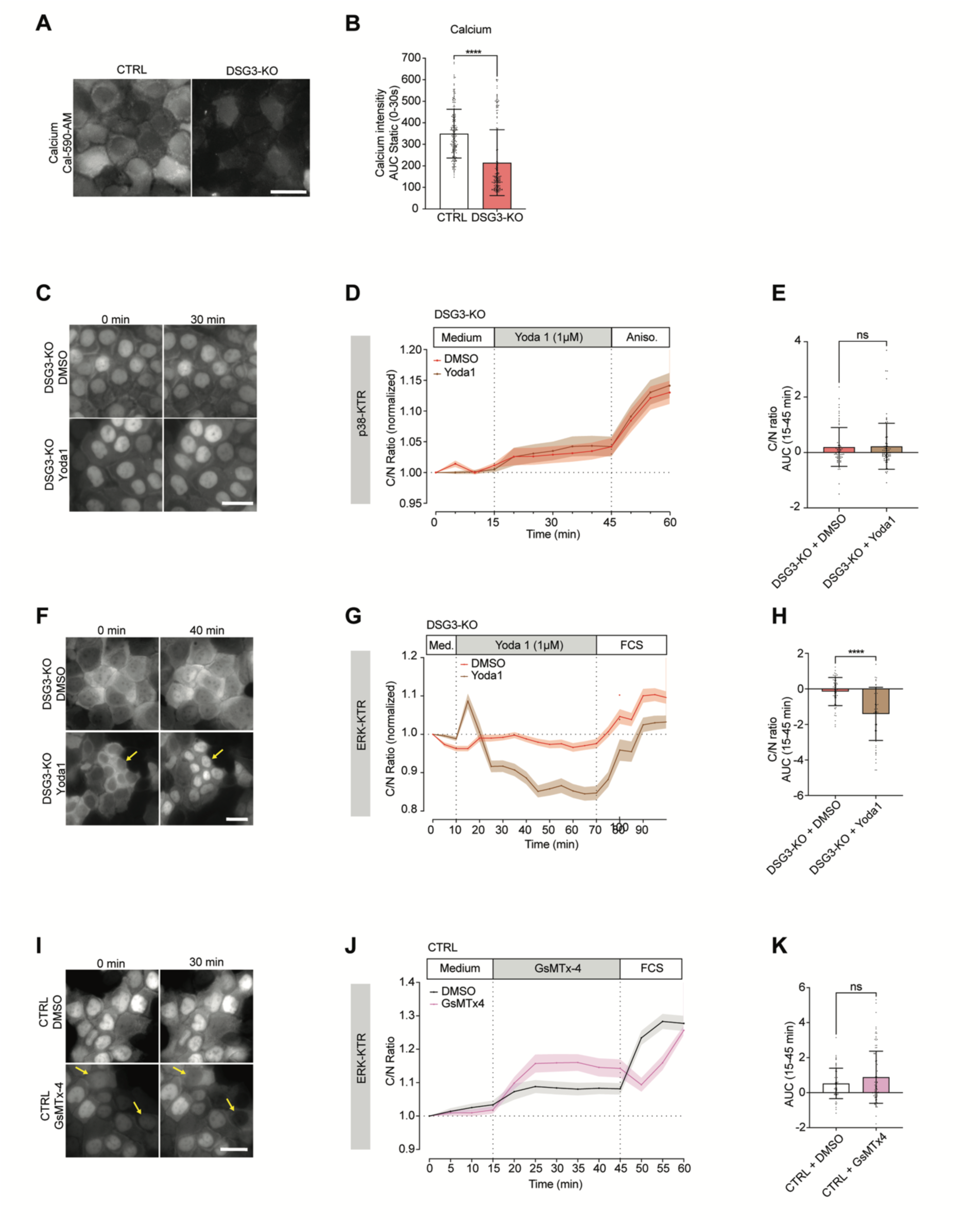
Piezo1 dampens ERK activity in static conditions. **A.** Representative images of intracellular calcium in CTRL and DSG3-KO cells stained with Cal-590-AM calcium indicator in static conditions. Scale bar 10 μm. **B.** Quantification area under the curve of calcium intensities imaged for 30 s in CTRL and DSG3-KO cells, N = 3, Mann-Whitney test. **C, F**. Time-lapse images of p38MAPK (C) and ERK (F) activity in DSG3-KO cells expressing the respective KTR sensors and treated with DMSO or Yoda1 in static conditions. **D, G.** Representation of p38MAPK (D) and ERK (G) dynamics depicted as the C/N ratio over time, normalized to the baseline. Data are represented as the mean ± SEM. **E, H**. The areas under the C/N ratio curve were calculated over the stimulation period, from 15 to 45 minutes, for p38MAPK (E) and from 10 to 70 minutes for ERK (H). Data represented as the mean ± SD. Individual cells from three independent experiments were analyzed using the Mann-Whitney test. **I.** Time-lapse images of ERK activity in CTRL cells treated with DMSO or GsMTx-4 in static conditions. **J.** Representation of ERK activity as C/N ratio variations over time in CTRL cells from (I). K. Quantification of the area under the curve in (J) calculated during the stimulation time from 15 to 45 minutes. Data represented as the mean ± SD. Individual cells from three independent experiments were analyzed using the Mann-Whitney test.

Interestingly, pharmacological Piezo1 activation by Yoda1 restored cell cohesion in DSG3-KO cells and also in those mutants which failed to increase adhesion (Fig. S4A, B). These results indicate that reduced PIEZO1 activity in DSG3-KO cells contributes to aberrant ERK signaling and show that adhesion depends on PIEZO1 function.

**Figure S4:**
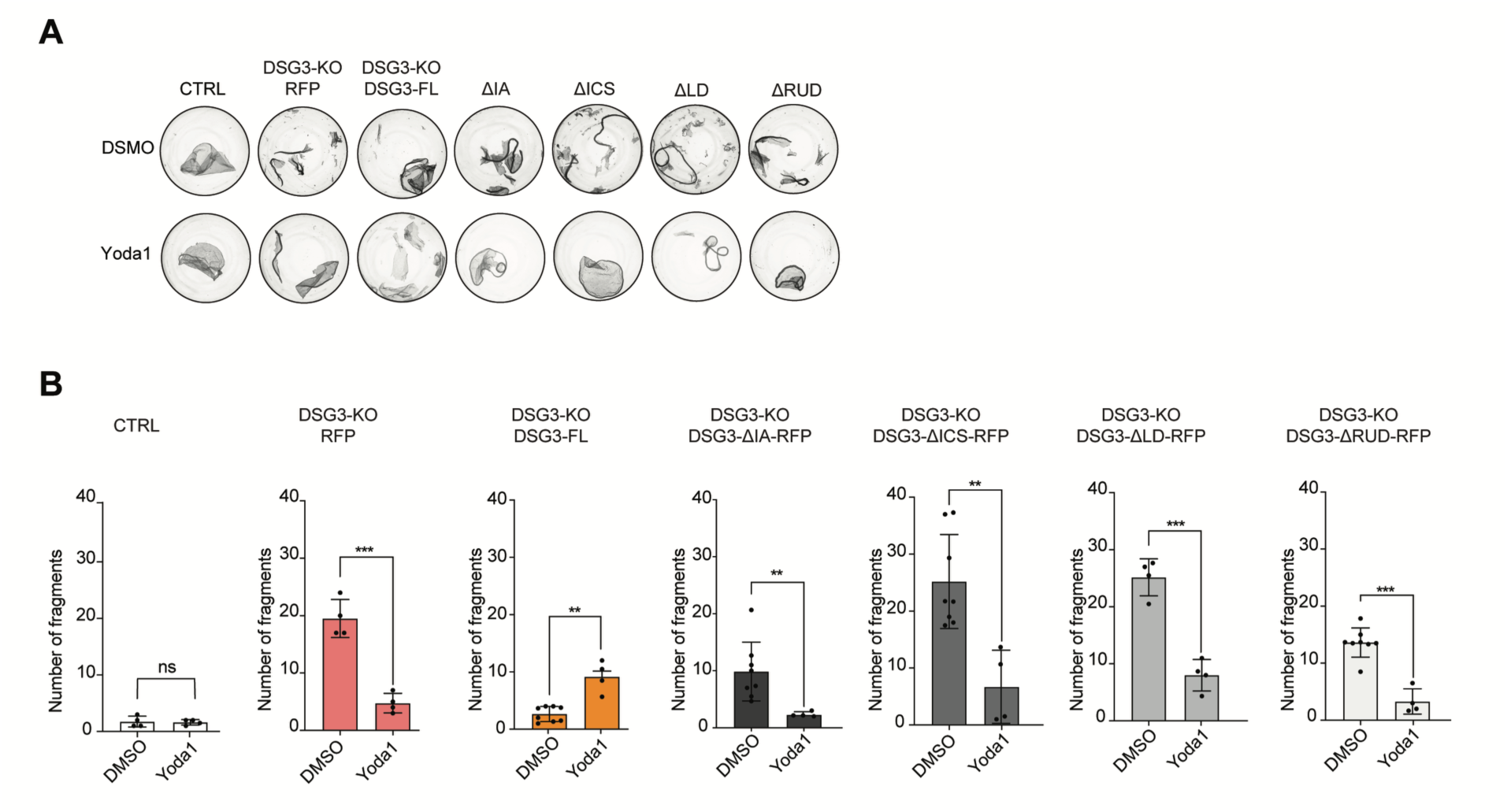
Piezo1 activity is important for cell-cell adhesion. **A**. Dispase-based dissociation assay of CTRL, DSG3-KO RFP, DSG3-FL, and DSG3 mutants treated with DSMO control or 0.5 µM Yoda1 for 24h. Representative images of cell monolayers after shearing are shown. **B**. Quantification of the number of fragments generated after shearing, N=4, Welch’s t test.

### DSG3 and Piezo 1 are required for p38MAPK and ERK responses to shear

So far, we investigated the relationship of DSG3 and MAPK signaling in static conditions. To approach mechanical stress present in the epidermis in a setting applicable to live cell imaging, keratinocytes were exposed to flow-induced shear stress. DSG3-KO and CTRL cells expressing the p38-KTR and ERK-KTR sensors were subjected to laminar flow (1 dyne/cm^2^) (Fig.S5A). Flow induced rapid and sustained p38MAPK activation in CTRL cells (Fig.4A-C, Fig.S5B, video 1). Interestingly, p38MAPK activation was absent in DSG3-KO cells (video 2). The ability of DSG3-KO cells to further increase p38MAPK and ERK activation was confirmed using anisomycin and fetal calf serum (FCS) as maximum activation control (Fig. S6A-F). Reconstitution with DSG3-FL partially recovered p38MAPK response, whereas all mutants failed to do so, suggesting that all DSG3 intracellular domains are required for modulating p38MAPK activity consistent with our observations under static conditions (Fig.4-A-C). Shear stress also elicited a rapid activation of ERK in CTRL cells, reaching a maximum by 20 min flow exposure (Fig.4D-F, Fig.S5C, video 3), in line with previous observations in endothelial cells (39). Again, loss of DSG3 markedly prevented ERK response to shear stress even resulting in a decreased activity over time (Fig.4D-F,video 4). Whereas re-expression of DSG3-FL restored ERK activation, none of the mutants increased ERK activity above the static baseline, except DSG3-ΔRUD, which showed partial ERK activation upon shear. These data demonstrate that DSG3 is needed for MAPK responses to shear. Further, all intracellular domains are required to elicit changes both in p38MAPK and ERK activity.

**Figure 4:**
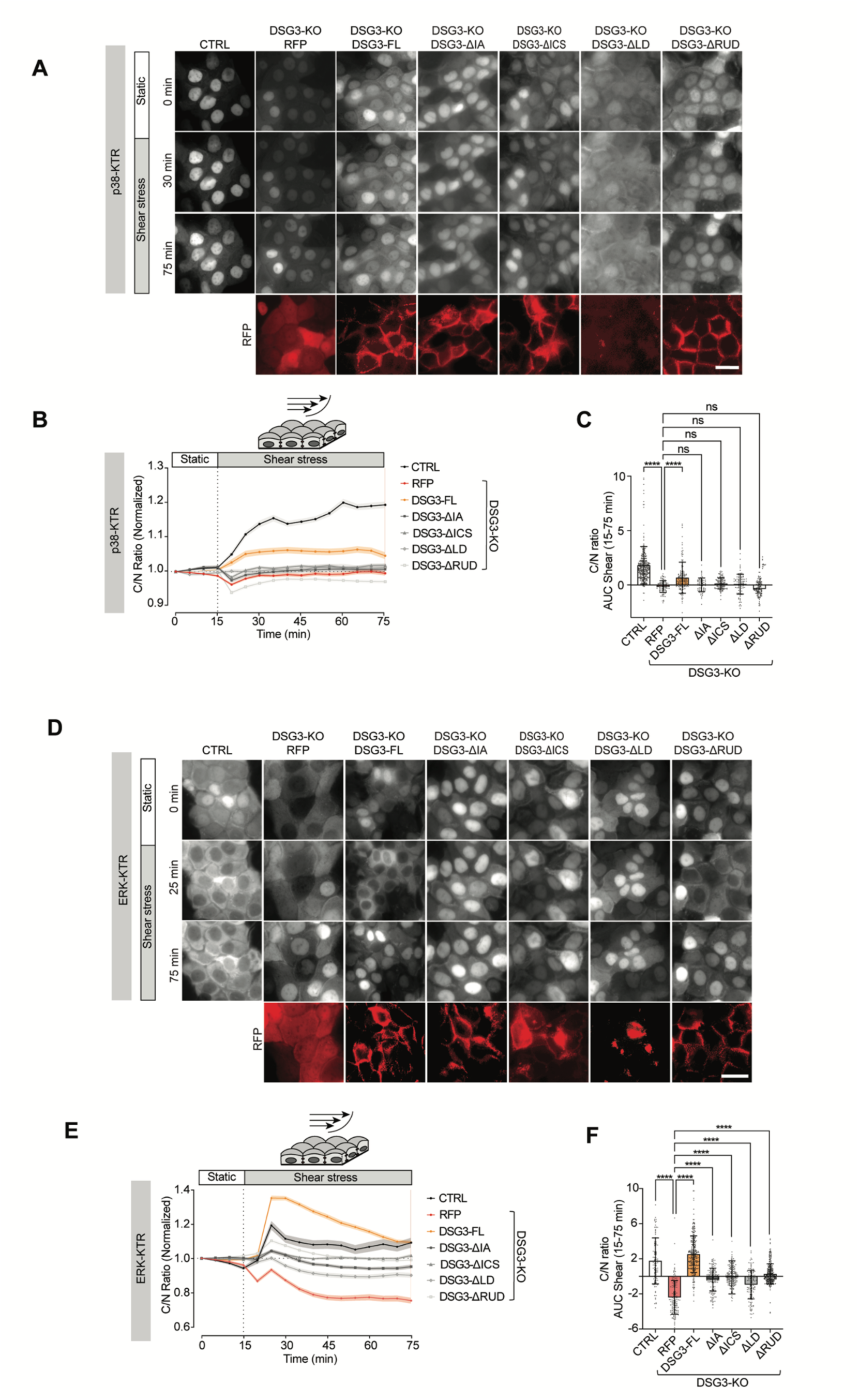
DSG3 is required to trigger p38MAPK and ERK activation upon shear stress. **A and D.** Time-lapse images of p38MAPK (A) and ERK (D) activity in CTRL, DSG3-KO, DSG3-KO DSG3-FL rescue, and the DSG3 mutant cells subjected to 1 dyne/cm^2^ flow shear stress. **B and E**. Representations of p38MAPK (B) and ERK (E) dynamics, depicted as the C/N ratio variations over time, normalized to baseline. **C and F**. The areas under the curve of the C/N ratio were calculated during shear stress between 15 and 75 minutes for p38MAPK and ERK in (B) and (E), respectively. One-way ANOVA Dunn’s multiple comparison test, data are represented as the mean ± SD from 99-295 cells of four independent experiments.

Application of shear stress induced rapid Ca^2+^ influx and sustained elevation of intracellular Ca^2+^ levels in a similar timescale as the MAPK responses (Fig. S7A-C). However, similar to MAPK activity, the Ca^2+^ influx was compromised in DSG3-KO cells.

**Figure S5:**
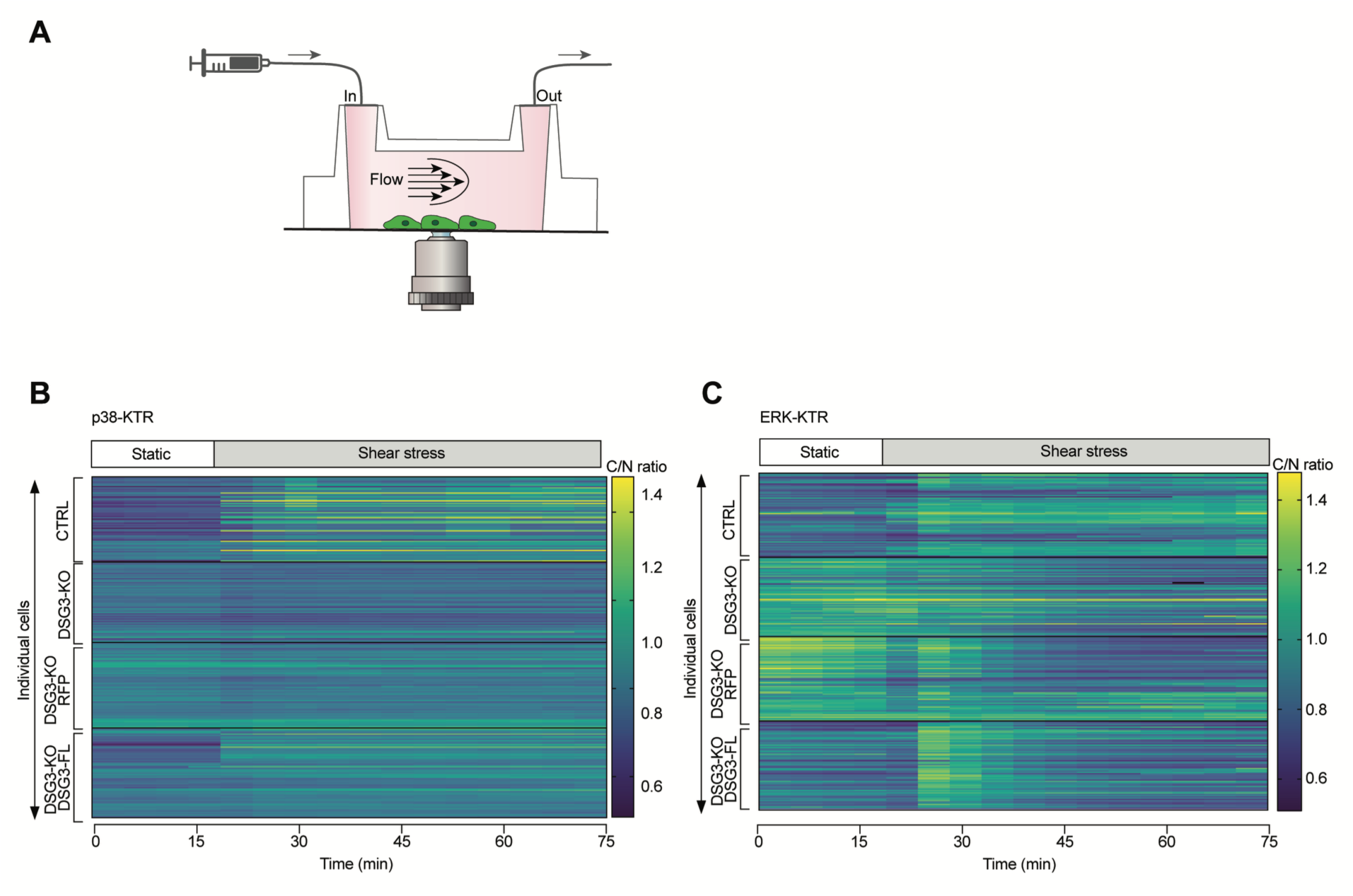
MAPK activation to shear stress in keratinocytes. **A.** Schematic of the microfluidic channel setup. HaCaT keratinocytes expressing the KTR sensor were cultured in the channels. When the cells reach confluence, a laminar-flow shear stress of 1 dyne/cm^2^ is applied while kinase dynamics are monitored by fluorescence live-cell imaging. **B-C**. Heatmap illustrating changes in C/N ratio over time from CTRL, DSG3-KO, DSG3-KO RFP, and DSG3-KO DSG3-FL keratinocytes expressing the p38-KTR (B) or the ERK-KTR (C) sensor, in response to shear stress. Each line represents one cell.

Importantly, activation of Piezo1 with Yoda1 restored p38MAPK activation in DSG3-KO cells subjected to shear stress (Fig. 5A-C). Similarly, activation of Piezo1 rescued ERK activation upon shear stress in the DSG3-KO cells (Fig.5D-F) while Piezo1 inhibition with GsMTx-4 prevented ERK activation in CTRL cells exposed to shear stress (Fig.5G-I). Together, these results demonstrate that Piezo1 activation and Ca^2+^ influx are required for both p38MAPK and ERK responses to shear stress.

**Figure 5:**
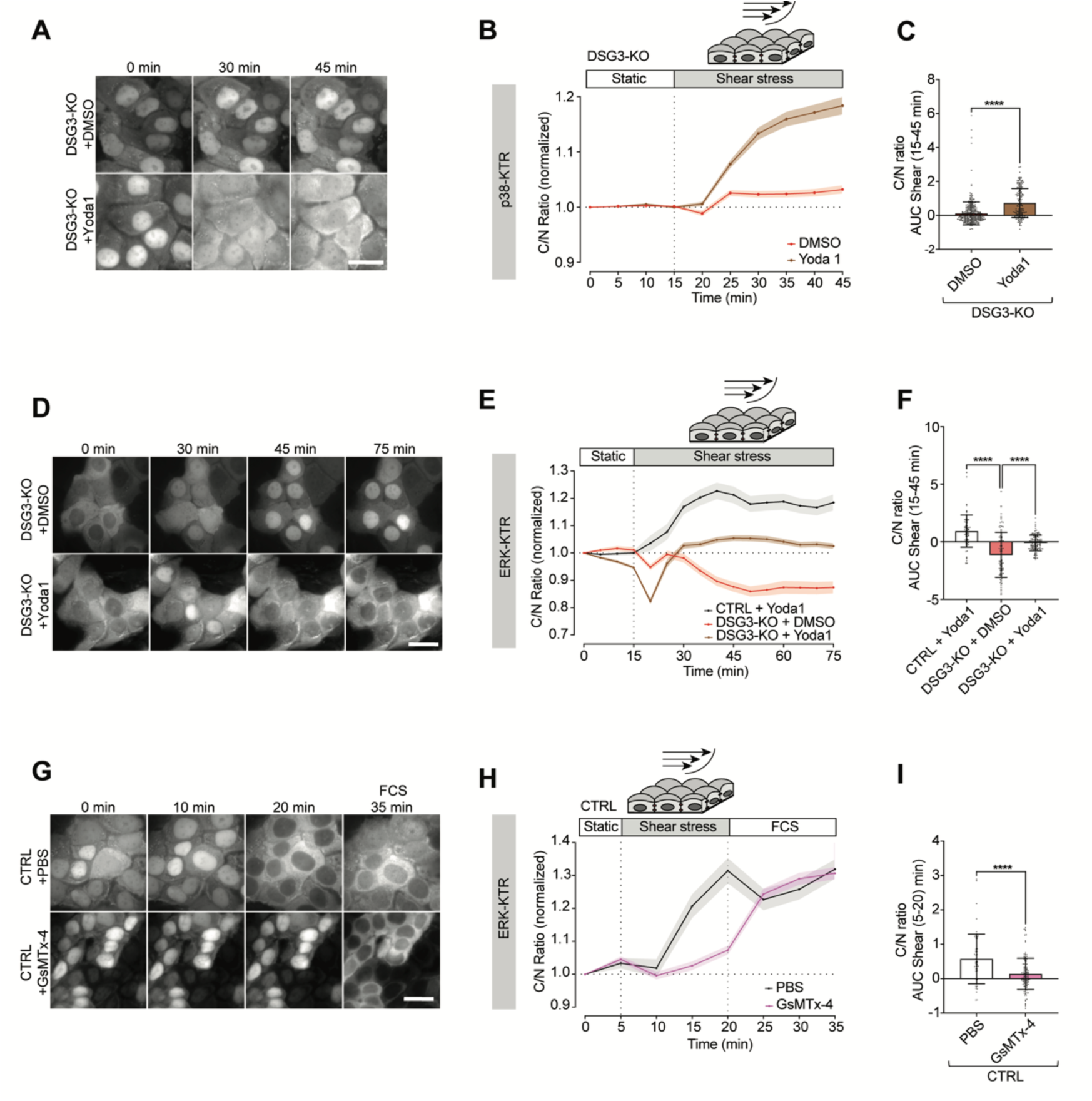
Activation of Piezo1 is essential for MAPK activation upon shear stress. **A and D**. Time-lapse images of DSG3-KO cells expressing the p38-KTR (A) and ERK-KTR (D) sensors. 1 dyne/cm^2^ shear stress was applied with DMSO or Yoda1-containing imaging medium. **B and E.** Representation of C/N ratio variations over time in response to shear stress, normalized to the baseline for p38MAPK (B) and ERK (E) activity. **C and F**. Quantification of the area under the curve of the C/N ratio in shear stress conditions (15-45min) in DSG3-KO cells treated with DMSO or Yoda1. For the p38-KTR experiment (C), 200 cells from three independent experiments were analyzed using the Mann-Whitney test. For the ERK-KTR experiment, 78-138 cells from three independent experiments were analyzed using one-way ANOVA with Dunn’s test. **G**. Representative images of CTRL cells expressing the ERK-KTR sensor subjected to shear stress and GsMTx-4, at the 20-minute timepoint, FCS (15%) was added as a control. **H**. Representation of C/N ratio variations over time in response to shear stress, GsMTX-4, and FCS, normalized to the baseline. **I**. Quantification of the area under the curve in shear stress conditions (5-20 min) in CTRL cells treated with PBS control or GsMTx-4, 138 cells from four independent exeriments were analyzed using the Mann-Whitney test.

## Discussion

In this study, we combined flow-induced shear stress with KTR-based live-cell imaging to investigate how keratinocytes regulate ERK and p38MAPK signaling both in static conditions and in response to mechanical forces. Using a DSG3 knockout model together with domain-specific DSG3 mutants, we identified DSG3 as a key regulator of steady-state kinase activity and responses to shear stress.

### DSG3-dependent MAPK regulation in static conditions

Deletion of DSG3 resulted in a reduced number of desmosomal proteins at the membrane, cytoskeletal changes, and weakened cell-cell adhesion. Comparison of CTRL and DSG3-KO proteomic profiles revealed proteins involved in force transmission and cytoskeletal regulation, including myosin-10, which modulates intracellular tensile forces (40), as well as integrins and LINC complex members. Pathway analysis identified changes in pathways linked to cellular stress responses and the RHOG GTPase cycle, which modulate the cytoskeleton, signaling, and mechanotransduction (41). Together, these findings indicate that DSG3 loss disrupts a network of junctional and cytoskeletal proteins required for both adhesion and mechanical force integration. This phenotype resembles the one observed in the blistering skin disease pemphigus vulgaris, in which autoantibodies against DSG3 impair Ca^2+^ fluxes, induce DSG3 internalization, and lead to cytoskeletal uncoupling and MAPK activation (42, 43). The similarity of these alterations to our results from knockout studies suggests that these mechanisms are largely caused by internalization or depletion of DSG3. Indeed, the pemphigus vulgaris phenotype can be prevented by overexpression of DSG3 in vitro (44), and activation of DSG3 transcription in vivo (45).

Unlike earlier WB-based methods for endpoint measurements of kinase activity, we employed real-time kinase monitoring using KTR sensors, which offer a clear advantage for elucidating signaling kinetics. The KTR sensors have been shown to be reliable tools for studying kinase activity across different cell types and for providing high-sensitivity measurements in living single cells (34, 35). In static conditions, DSG3 deletion increased basal p38MAPK and ERK activity, demonstrating a dependence on DSG3 in dampening MAPK signaling in the steady state. Importantly, all DSG3 mutants, even the cytosolically localized DSG3-ΔLD mutant, were able to restore low basal ERK activity. This indicates that membrane localization of DSG3 is required to suppress ERK signaling under static conditions. With regard to cell adhesion, none of the mutants was able to reestablish full cell cohesion. This outlines that cell adhesion and ERK suppression is at least partially uncoupled. This is a relevant observation, as a broader concept in the field suggests that loss of desmosomal cadherin binding leads to changes in intracellular signaling (43). It is still possible that interactions of individual DSG3 extracellular domains, for example, in cis with other Dsg3 molecules, are sufficient to prevent ERK signaling under conditions of impaired intercellular adhesion.

In contrast to ERK, none of the DSG3 mutants was able to restore basal p38MAPK activity, indicating that the regulation depends on the integrity of the entire DSG3 intracellular tail. Again, p38MAPK suppression was partially uncoupled from cell adhesion, because the IA and RUD mutants ameliorated the loss of cell adhesion but had no effect on p38MAPK activity.

One effector of DSG3-dependent signaling appears to be Piezo1. Expression of Piezo1 was reduced under static conditions upon DSG3 loss, and exogenous activation of this channel restored decreased ERK activity. In contrast, modulating Piezo1 activity had no effect on p38MAPK, again indicating that these two kinases are regulated separately.

With regard to p38MAPK, we have previously shown that p38MAPK can interact with DSG3, which may indicate why the intracellular domains are required for the modulation. Alternatively, p38MAPK activity depends on the balance between phosphorylation by upstream activators and dephosphorylation by negative regulators such as phosphatases (DUSPs). Indirectly, DSG3 may suppress p38MAPK by interfering with upstream pathways such as EGFR/Src and Ca^2+^/PKC, mechanisms that have been shown to be relevant for p38MAPK activation in pemphigus (12, 22, 46, 47).

### DSG3-dependent MAPK regulation under dynamic conditions

Using shear stress as proxy for mechanical perturbation, we observed an activation p38MAPK and ERK in keratinocytes, consistent with observations in cardiomyocytes and endothelial cells (48). Notably, we show that DSG3 plays a dual role in regulating MAPK activity, which differs between static and shear-stress conditions. Under static conditions, DSG3 is required to dampen p38MAPK and ERK activity, whereas in response to shear, it appears to be required for MAPK activation. We cannot fully exclude the possibility that the “pre-activation” of p38MAPK and ERK under static conditions caused by the knockout of DSG3 interferes with the response to the onset of shear. However, application of positive controls resulted in further increased MAPK activation in DSG3-KO cells, indicating that DSG3-KO cells are not already maximally activated.

Interestingly, DSG3 has been shown to integrate into two distinct pools with different cytoskeletal components, one interacting with actin on the cell surface and the other with intermediate filaments at cell-cell contacts (49). This dual cytoskeletal anchoring mechanism may be related the distinct regulation of signaling by DSG3. It is possible that an actin-associated extra-desmosomal DSG3 pool was responsible for the shear response, whereas desmosomal DSG3 modulates MAPK activity under steady-states.

Re-expression of full-length DSG3 partially restored p38MAPK activation and fully restored ERK activation in response to shear, although dynamics were altered. Similar to the static condition, none of the DSG3 mutants restored p38MAPK activation upon shear stress. This demonstrates that full DSG3 tail integrity is also important for the rapid p38MAPK responses to shear. Regarding ERK activity, only the DSG3-ΔRUD mutant showed partial ERK activation under shear, suggesting that the RUD domain is dispensable for the ERK response.

MAPK activation by shear stress is well established in endothelial cells and is mediated by a junctional mechanosensory complex consisting of PECAM-1 and VE-cadherin, which leads to ligand-independent activation of VEGFR2 (50, 51). In parallel, shear stress activates Piezo1, leading to calcium influx, triggering activation of Src-family kinases and non-canonical EGFR phosphorylation (52–55). Together, these pathways converge to activate the Ras/Raf/MEK/ERK cascade (53, 56). The crosstalk between these junctional complexes and ion channels contributes to a mechanotransduction machinery that regulates endothelial adaptation to flow (52, 53, 56–58). Although keratinocytes are not exposed to constant fluid flow, the mechanisms of interaction between junctional complexes and Piezo1 could also apply to keratinocytes during physiological mechanical challenges.

Consistently, shear stress induced a calcium influx in CTRL cells, with an initial sharp, transient increase followed by a gradual secondary increase. DSG3 deletion altered calcium kinetics, resulting in a smaller initial peak and specifically impacting the second phase. This could suggest a disruption in store-operated Ca^2+^ entry or IP3-mediated calcium release from the endoplasmic reticulum (59–61). Interestingly, recent studies have shown that the ER is physically linked to desmosomes and that calcium release via IP3R signaling directly affects desmosome stability (62–65), suggesting a potential ER-IP3R dysfunction in the DSG3-KO that would be interesting to decipher in further studies. Moreover, calcium signals are known to be mediated by Ca^2+^ -binding proteins such as calmodulin kinase (CaMK) or PKC, which are upstream activators of MAPK (66).

Although it is unclear whether the altered Ca^2+^ dynamics in DSG3 KO cells are a result of impaired Piezo1 function, our data demonstrate that dynamic MAPK responses depend on Piezo1. Pharmacological activation of Piezo1 restored both ERK and p38MAPK activation under shear stress in DSG3-KO cells, whereas Piezo1 inhibition dampened ERK activation in CTRL cells. So far, it is unclear how DSG3 and Piezo1 responses are connected. Interestingly, Bio-ID studies did not indicate that Piezo1 interacts with desmosomal cadherins (67, 68). Thus, Piezo1 is likely indirectly stabilized by DSG3. Piezo1 activity is regulated mainly by membrane curvature, but also by direct phosphorylation or cytoskeletal coupling involving cortical actin and myosin that transmit internal forces (69, 70). DSG3 may regulate Piezo1 activity indirectly by modulating the actin cytoskeleton and loss of DSG3 may affect Piezo1 mechanosensitivity. Piezo1 gating is known to depend on cortical actin stiffness (71–74), which may be altered in DSG3-KO cells due to cytoskeletal disruption. AFM-based studies showed that altered cortical actin reduces cellular stiffness (75–78). Thus, DSG3 may regulate Piezo1 activity indirectly by stabilizing the actin cytoskeleton rather than by controlling Piezo1 expression alone. Despite their lack of direct interaction, desmosomes and Piezo1 are both localized to lipid rafts in the membrane (79–81). Piezo1, in turn, is known to interact with EGFR and activate ERK and p38MAPK signaling (54, 56, 82). Together, these findings are consistent with a role for DSG3 in indirectly modulating Piezo1 activity and Ca^2+^ dynamics, which are required for MAPK signaling responses.

## Conclusion

Together, our data support a mechanistic model for how DSG3 regulates p38MAPK and ERK signaling (Fig. 6). In static conditions, DSG3 inhibits p38MAPK activity, thereby stabilizing keratin filaments and intercellular adhesion. In parallel, DSG3 stabilizes Piezo1 expression and its basal activity, thereby dampening ERK signaling and contributing to cell cohesion. In response to mechanical forces, DSG3 ensures stable desmosomes and a rigid cortical actin cytoskeleton, thereby maintaining Piezo1 gating under mechanical stress, where proper Piezo1-mediated calcium influx is important for ERK and p38MAPK activation. Together, our study reveals DSG3 as a signaling hub that integrates external mechanical signals, intracellular pathways, and cytoskeletal modifications to help the cell adapt to its environment.

**Figure 6:**
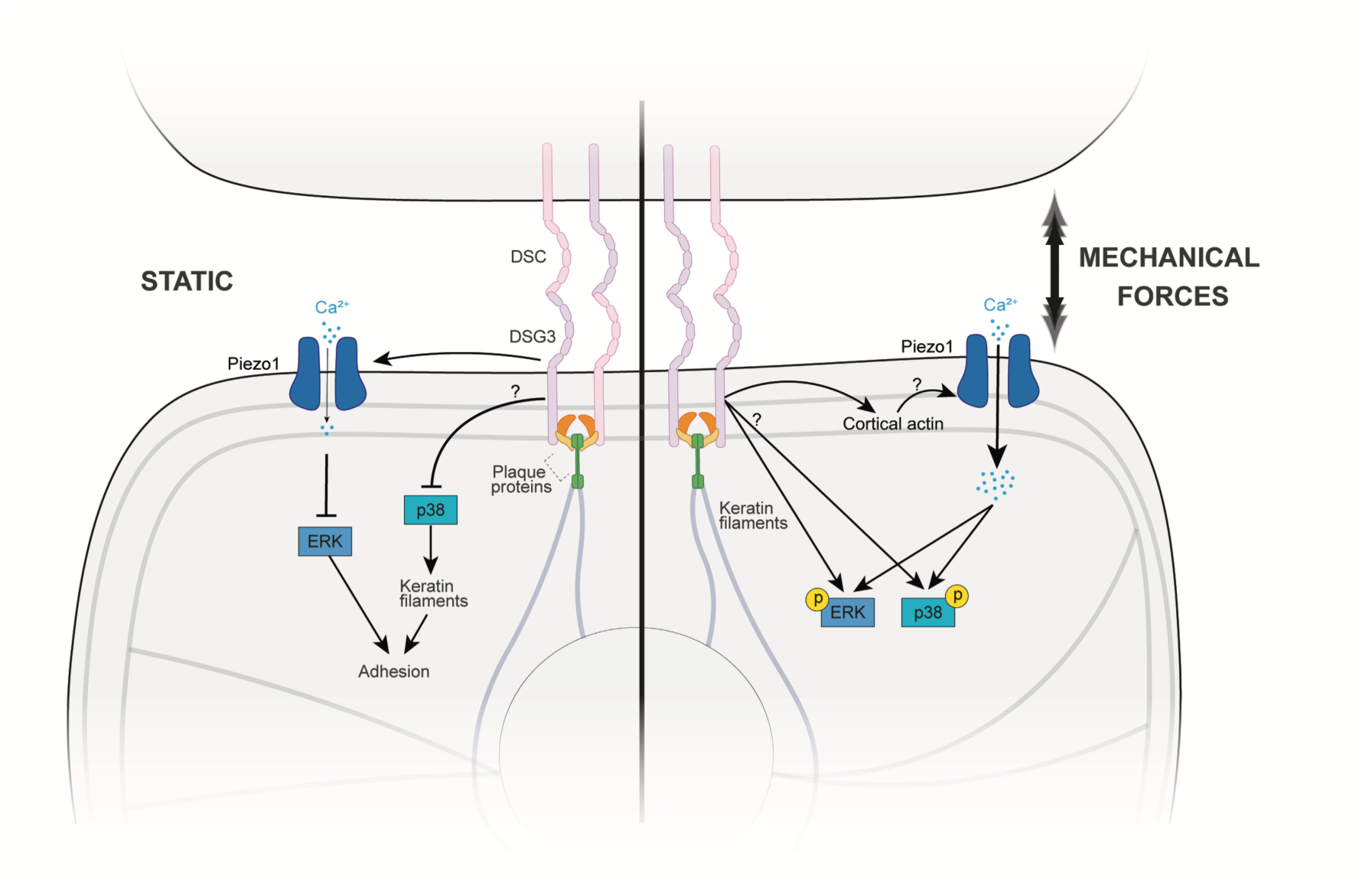
Working model: Schematic of the proposed DSG3-dependent modulation of MAPK activity in static and dynamic conditions. Arrows indicate activation, while T-bars represent inhibition. In static: DSG3 stabilizes Piezo1 expression and maintains its basal activity, thereby dampening ERK activity. DSG3 also dampens p38MAPK activity. Together, DSG3 maintains a signaling hub that dampens MAPK activity, thus stabilizing keratin and cortical actin filaments and contributing to strong intercellular adhesion. In shear stress conditions, Piezo1 activation triggers a significant influx of calcium, which in turn activates p38MAPK and ERK.

## Material and Methods

### HaCaT cell culture

The spontaneously immortalized HaCaT (Human adult high Calcium low Temperature) keratinocyte cell line was cultured in a humidified atmosphere of 5% CO2 and 37 °C in Dulbecco’s Modified Eagle Medium (DMEM) (Sigma-Aldrich, D6546) containing 1.8 mM Ca^2+^ and complemented with 10% fetal bovine serum (Merck, S0615), 50 U/mL penicillin (VWR, A1837.0025), 50 μg/mL streptomycin (VWR, A1852.0100) and 4 mM L-glutamine (Sigma-Aldrich, G7513).

### Cloning of sgRNA

The oligos for sgDSG3 and sgNT1 (Table 1) were synthesized by Microsynth using the T4 PNK enzyme (NEB, #0201S). The oligos were then annealed and ligated into Esp3I-digested lentiCRISPR_v2 (Addgene, #52961) using T4 ligase (NEB, #M0202S). The ligated product was transformed into competent DH5α bacteria and plated onto agar plates containing Ampicillin (100 μg/ml). Picked colonies were then amplified in a 3 ml medium containing ampicillin (100 μg/ml). Plasmids were isolated using a Miniprep kit (Macherey-Nagel, #300287) and sequenced using the U6 forward primer.

### Cloning of murine DSG3 mutants

The PCR of the mDSG3 plasmid was performed using the Platinum SuperFi II DNA polymerase (Fisher Scientific, product number 16410771), according to the manufacturer’s protocol. The primers used, including an overhang for the next domain to amplify the targeted domains, are listed in Table 2.

The annealing temperature for the primers was set to 56 °C. The PCR products were purified using a PCR and Gel Purification Kit (Qiagen, #28506). To fuse the amplified domains of mDSG3, the necessary PCR products (1.25 ng each) were amplified in a single PCR reaction for 15 cycles (initial denaturation at 95 °C for 2 minutes, followed by 15 denaturations at 95 °C for 30 seconds, annealing at 56 °C for 30 seconds, and extension at 72 °C for 45 seconds, with a final extension at 4 °C forever). Then, Asc1_mDSG3_for and mDSG3_Not1_rev (final concentration of 0.5 µM), as well as 0.5 μL of Platinum Super Fi II DNA polymerase, were added and amplified for an additional 20 cycles (initial denaturation at 95 °C for 2 minutes, followed by 20 denaturations at 95 °C for 30 seconds, annealing at 56 °C for 30 seconds, and extension at 72 °C for 45 seconds, with a final extension at 4 °C forever). The PCR product was purified using a PCR and Gel Purification Kit (Qiagen, #28506) and digested with Not1 (Bioconcept, #R3189L) and Asc1 (Bioconcept, #R0558L) for three hours at 37 °C. The digested product was loaded onto a 1% agarose gel and purified from the gel using a PCR and Gel Purification Kit (Qiagen, #28506). The insert was ligated into the Asc1/Not1-digested pLenti-C-mRFP-P2A-Puro vector at a ratio of 3:1 overnight at 16 °C using T4 ligase (Bioconcept, #300361). Positive clones were sequenced using the following primers: pLenti_RFP_V2_for_seq (AGCAGAGCTCGTTTAGTGAACC) and pLenti_RFP_rev_seq (TGGTGGTTGTTCACGGTGC).

### Generation of lentiviral constructs and stables cell lines

Kinase Translocation Reporter plasmids were purchased from Addgene: pLentiCMV Puro DEST ERKKTRClover (#59150) for ERK and pLentiPGK Puro DEST p38KTRClover (#59152) for p38MAPK. Stbl3 bacterial strains were plated on LB agar containing ampicillin (100μg/mL), and single colonies were amplified in 2 mL LB medium with ampicillin (100 μg/mL). Plasmids were purified using NucleoBond Xtra Midiprep kit (Macherey-Nagel, #740410.50) and verified by sequencing using the primers indicated in Table 3.

Lentiviral particles were generated according to standard procedures. HEK293T cells were transfected with lentiviral packaging vector psPAX2 (Addgene, #12259), the envelope vector pMD2.G (Addgene, #12260), and the respective construct plasmid using TurboFect (Thermo Fisher Scientific). 48 h after transfection, virus-containing supernatant was collected and concentrated using Lenti-Concentrator (OriGene) for 2h at 4°C. HaCaT keratinocytes were stably transduced with the respective virus particles in an equal ratio using 5 µg/ml polybrene (Sigma-Aldrich) according to the manufacturer’s instructions. 24h after transduction, medium was exchanged and puromycin 1 μg/mL was added in the culture media for drug selection. Cells were cultivated for at least 1 week under selection then imaged to confirm fluorescent protein expression before starting with the respective experiments. Cells expressing both the KTR sensor and RFP-tagged DSG3 mutants were sorted for GFP⁺/RFP⁺ double-positive cells using fluorescence-activated cell sorting (FACS).

### Western blot

HaCaTs were seeded in 24-well plates until reaching confluency. Cell monolayers were lysed with SDS lysis buffer (25 mM HEPES, 2 mM EDTA, 25 mM NaF, 1% SDS, pH 7.6) supplemented with an equal volume of a protease inhibitor cocktail (cOmplete, Roche Diagnostics) and phosphatase inhibitor (PhosSTOP™, Roche). Lysates were sonicated and the total protein amount was determined with a BCA protein assay kit (Thermo Fisher Scientific) according to the manufacturer’s instructions. The proteins were denatured by heating in Laemmli buffer, for 10 minutes at 95°C. After run, membranes were blocked in Odyssey blocking buffer (Li-Cor) for 1 hour at room temperature (RT). The following primary antibodies were diluted with Odyssey blocking buffer in TBS containing 0.1% Tween 20 (Thermo Fisher Scientific) and incubated overnight at 4°C, with rotation: mouse GAPDH mAb (Santa Cruz Biotechnology, #sc-47724), rabbit phospho-p38MAPK Thr180/Tyr182 (Cell Signaling Technology, #4511S), rabbit p38MAPK (Cell Signaling Technology, #9212S), rabbit ERK 1/2 (p44/42) (Cell Signaling Technology, #9102), phospho-ERK (Santa Cruz, sc-7383), mouse plakoglobin (Progen, #61005), mouse desmoplakin 1/2 (Progen, #61003), rabbit Piezo1 (Proteintech, # 15939-1-AP), rabbit Desmoglein-3 (5G11, Santa Cruz Biotechnology, # sc-53487), Desmoglein-2 (10G11, Acris OriGene, #BM5016), mouse Pan Cytokeratin mAb (Thermo Fisher, #41-9003-82), Phalloidin (Thermo Fisher, #21833). Goat anti-mouse 800CW and goat anti-rabbit 680RD (Li-Cor, #925-32210 and 925-68071) were used as secondary antibodies, incubated for 1 hour at RT. Odyssey FC imaging system was used for imaging the blots, and band intensity was quantified with Image Studio (both Li-Cor).

### Shear stress live-cell Imaging experiments

Cells were seeded at 3 x 10^4^ cells per channel onto IBIDI μSlide VI 0.4 flow chambers (IBIDI, ibiTreat, #80606) and cultured with daily medium changes. At 90% confluency, cells were serum-starved overnight in medium containing 1% FBS. On the day of the experiment, cells were incubated with 50 ng/mL Hoechst 33342 (Fisher Scientific, #62249) for 30 min at 37°C to facilitate nuclear segmentation. Cells were then switched to imaging medium composed of DMEM without phenol red (Sigma-Aldrich, D1145) supplemented with 1% FBS (Merck, S0615), 50 U/mL penicillin (VWR, A1837.0025, D6546), 50 μg/mL streptomycin (VWR, A1852.0100), 4 mM L-glutamine (Sigma-Aldrich, #G7513), 10 mM sodium pyruvate (Merck, #1.06619.0050) and 25mM Hepes (Sigma-Aldrich, #H3375) for at least 1h prior to imaging. Temperature was maintained at 37°C throughout experiments. Flow shear stress (1 dyne/cm²) was applied for the indicated duration using a syringe pump (Harvard Apparatus, Pump 11 Elite Infuse Only, #704500) and a 60 ml syringe (B. Braun Omnifix #4617509F). Images were acquired every 5 min. Imaging was performed with an IX83 inverted fluorescence microscope (Olympus) equipped with a 60x oil immersion objective (PlanApo N 60x/1.42 oil, Olympus) and an Orca-Flash 4.0 V3 digital CMOS camera (Hamamatsu, #C13440-20CU) controlled by the CellSens Imaging software (Olympus).

### Image analysis of the KTR

Nuclei were segmented based on Hoechst staining, and a 5-pixel cytoplasmic ring was applied to quantify cytoplasmic signal using eDetect under MATLAB (R2025a, MathWorks). Tracking and manual correction were performed as previously described (Han et al., 2018). Median intensity values were used to calculate cytoplasmic-to-nuclear (C/N) ratios, reflecting kinase activity (workflow detailed in Fig. S2B). Time-course data were normalized to each cell’s basal value. The area under the curve (AUC) was calculated as the sum of all normalized values minus the number of time points, representing cumulative deviation from baseline. Population data are presented as mean ± SEM from pooled single-cell measurements across independent experiments. Single-cell traces in heatmaps are provided in the Supplementary Materials (Fig.S5B-C, S6C-F).

### Calcium Imaging

Cells were loaded with 4 µM Cal-590 AM (Aat Bioquest, #20510) in imaging buffer (modified HBSS; Sigma H8264 supplemented with 2 mM CaCl₂·2H₂O, 2 mM MgCl₂·6H₂O and 10 mM HEPES, pH 7.4) for 60 min at 37 °C. The measurement buffer contained (in mM): 137 NaCl, 5.37 KCl, 2 CaCl₂, 2 MgCl₂, 0.34 Na₂HPO₄, 0.44 KH₂PO₄, 4.17 NaHCO₃, 5.55 D-Glucose, and 10 HEPES. Following dye loading, fresh measurement buffer was added before the experiment. To serve as a negative control, imaging buffer containing 5 mM EDTA was used to chelate extracellular Ca²⁺. Intracellular Ca²⁺ dynamics were quantified as changes in fluorescence intensity over time.

### Dispase-based dissociation assay

HaCaT cells were seeded at 100 000 cells per well in 24-well plates (Greiner Bio-One) and cultured under standard conditions (37°C, 5% CO₂) to form a confluent monolayer and treated for 24h with SB202190 30μM (Sigma Millipore, #559388), UO126 10μM (Sigma-Aldrich, #19-147) or Yoda1 0.5μM (Sigma-Aldrich, #SML1558). Prior to the assay, monolayers were washed thoroughly with Phosphate-buffered saline (PBS). The cell monolayer was then incubated with dispase II (2.4 U/mL; Sigma-Aldrich) for 20 minutes at 37°C to detach the intact cell monolayers from the well bottom. After incubation, the dispase II solution was carefully removed and replaced with HBSS (Sigma-Aldrich, #H8264-6X1L). Mechanical stress was subsequently applied by pipetting the detached monolayer 20 times using an electrical pipet set to 350 µl (Xplorer 1000, Eppendorf, #L49475G). The fragments were documented and quantified with a stereo microscope (Olympus, #SZX2-TR30) with an attached camera (Canon, #EOS 800D). Each experimental condition was performed in triplicate wells, and the entire experiment was repeated at least three independent times.

### Immunostaining

Cells were grown on 13-mm glass coverslips and fixed with either 2% PFA (Thermo Fisher Scientific) in PBS at room temperature or ice-cold methanol (Merck Millipore) for 10 min on ice. Cells were permeabilized with 0.1% Triton X-100 in PBS for 10 min and blocked with 3% bovine serum albumin (BSA, VWR, #422351S) and 1% normal goat serum (NGS, Jackson ImmunoResearch, #005-000-121) in PBS for 1 h in a humidified chamber. The following primary antibodies were incubated overnight at 4°C: mouse Dsg3 mAb (clone 5G11, Thermo Fisher, #32-6300), Phalloidin CruzFluor 488 (Santa Cruz, #sc-363791), pancytokeratin (AE1/AE3) efluor 570 (#41-9003-80; eBioscience), mouse desmoplakin 1/2 (Progen, #61003), rabbit Piezo1 (Proteintech, # 15939-1-AP), and Phalloidin (Thermo Fisher, #21833). Following primary antibody incubation, cells were washed three times with PBS and AlexaFluor (AF488, AF568) conjugated anti rabbit or anti-mouse antibodies (#A-11008, A-11004; Thermo Fisher Scientific) were added and incubated for 1h at room temperature. DAPI (Sigma-Aldrich) was added for 10 min for nuclei staining. Cells were washed three times with PBS and mounted with Prolong Diamond Antifade (Thermo Fisher Scientific). Image acquisitions were done using a Stellaris 8 Falcon confocal microscope (Leica) with an HC PL APO CS2 63×/1.40 oil objective. Image analysis was done with ImageJ software. DSP mean intensity at the cell membrane was quantified by drawing a region of interest (ROI) around the plasma membrane, and the DSP intensity was then normalized by the respective length of the plasma membrane. For Piezo1, mean intensity was measured in a rectangular ROI and cell nuclei were detected using DAPI. The mean intensity of the staining was divided by the number of nuclei. Keratin distribution was analyzed by drawing a fixed-size ROI rectangle at the cell–cell contact sites (10 microns) and measuring the mean plot profile intensities along this area for individual cells.

### Proteomics

Cells were lysed with buffer containing 100 mM TEAB pH=8.5 / 5% SDS / 10 mM TCEP by 10 min heating at 95° followed by 10 min sonication (30s on, 30s off per cycle) on a Pixul® system (Active Motif) using following settings: Pulse[N]: 50; PRF[kHz]: 1; burst rate[Hz]:20. Protein extracts were alkylated using 15 mM iodoacetamide at 25° C in the dark for 30 min. For each sample, 100 µg of protein lysate was captured, digested, and desalted using STRAP cartridges (Protifi). In brief, samples were acidified by addition of 10% TFA (1:10) and then S-trap buffer (90% methanol, 100 mM TEAB pH 7.1) was added to the samples (6:1). Samples were briefly vortexed and loaded onto S-trapTM micro spin-columns (Protifi) and centrifuged for 1 min at 4000 g. Flow-through was discarded and spin-columns were then washed 3 times with 150 µL of S-trap buffer (each time samples were centrifuged for 1 min at 4000 g and flow-through was removed). S-trap columns were then moved to the clean tubes and 20 µL of digestion buffer (50 mM TEAB pH 8.0) and trypsin (at 1:25 enzyme to protein ratio) were added to the samples. Digestion was allowed to proceed for 1h at 47 °C. After, 40 µL of digestion buffer was added to the samples and the peptides were collected by centrifugation at 4000 g for 1 minute. To increase the recovery, S-trap columns were washed with 40 µL of 0.2% formic acid in water (400g, 1 min) and 35 µL of 0.2% formic acid in 50% acetonitrile. Eluted peptides were dried under vacuum and stored at -20 °C until further processing.

Peptides were resuspended in 0.1% aqueous formic acid and 0.2 ug of peptides subjected to LC–MS/MS analysis using an Orbitrap Exploris 480 Mass Spectrometer fitted with an Vanquish Neo (both Thermo Fisher Scientific) and a custom-made column heater set to 60°C. Peptides were resolved using a RP-HPLC column (75μm × 30cm) packed in-house with C18 resin (ReproSil-Pur C18–AQ, 1.9 μm resin; Dr. Maisch GmbH) at a flow rate of 0.2 μLmin-1.

Separation gradient was: from 2% Buffer B to 10% Buffer B over 5 min, to 35% Buffer B over 45 min, to 50% Buffer B over 10 min. The mass spectrometer was operated in data-independent acquisition (DIA) mode. Each MS1 scan was acquired at a resolution of 120,000 FWHM (at 200 m/z) and a scan mass range from 390 to 910 m/z. Maximum injection time was set to auto mode with a normalized AGC target set to 300%. Each survey scan was followed by a set of DIA scans acquired at a resolution of 15,000 FWHM. Precursor mass range was set to 400-900 m/z, the isolation windows size to 10 m/z and the HCD normalized collision energy (NCE) to 28%. For MS2 scans, maximum injection time was set to 22 ms with a normalized AGC target set to 3000%. The acquired raw files were searched using SpectroNaut (v19.0, directDIA workflow, default settings) against a human database (downloaded from Uniprot on 20220222) using the following search criteria: full tryptic specificity was required (cleavage after lysine or arginine residues, unless followed by proline); 3 missed cleavages were allowed; carbamidomethylation (C) was set as fixed modification; oxidation (M), N-acetylation (N-term) were applied as variable modifications. The identified phospho peptides were exported as tsv files and analyzed using MSStats R package v.4.13 (83).

STRING database was used to identify molecular and biological pathways associated with the significantly modulated proteins across the samples. X-axis denotes −log10 of FDR values, with a cutoff set to 0.05 for significance.

## Statistics

Statistical analyses were performed using GraphPad Prism 8 (GraphPad Software). Data sets were first assessed for normality using the Shapiro-Wilk test. Comparisons between two groups were performed using two-tailed Student’s t tests, while comparisons involving more than two groups were analyzed by one-way or two-way ANOVA, as appropriate. P values < 0.05 were considered statistically significant. All data are presented as mean ± SD unless otherwise indicated. A minimum of three independent biological replicates was included for each experiment. Figures were prepared using Adobe Illustrator 29.8.3 (Adobe Inc.).

## Acknowledgements

The authors would like to acknowledge Pascal Lorentz (Microscopy Core Facility, University of Basel, Switzerland) for his valuable advice and support in microscopy.

The study was supported by the Swiss National Science Foundation (197764_1 to V. Spindler)

## Abbreviations

AFM: Atomic force microscopy
Ca^2+^: Calcium
CaMK: Calmodulin kinase
CRISPR/Cas9: Clustered Regularly Interspaced Short Palindromic Repeats and CRISPR-associated protein 9
CTRL: Control
DMSO: Dimethyl sulfoxide
DSG: Desmoglein
DSG1: Desmoglein 1
DSG2: Desmoglein 2
DSG3: Desmoglein 3
DSG3-KO: Desmoglein 3 knockout
DSP: Desmoplakin
DTD: Desmoglein terminal domain
DUSP: Dual specificity protein phosphatase
EGFR: Epidermal growth factor receptor
ER: Endoplasmic Reticulum
ERK: Extracellular Signal-Regulated Kinase
FCS: Foetal calf serum
FL: Full-length
FRET: Förster resonance energy transfer
GsMTx-4: Grammostola spatulata mechanotoxin 4
HaCaT: Human adult high Calcium low Temperature
IA: Intracellular anchor
ICAM1: Intercellular adhesion molecule 1
ICS: Intracellular cadherin-like sequence
IP3: Inositol trisphosphate
ITGB6: Integrin beta-6
JUP: Junction plakoglobin
KTR: Kinase translocation reporter
LD: Linker domain
LINC: Linker of Nucleoskeleton and Cytoskeleton
MAP2K2: Dual specificity mitogen-activated protein kinase kinase 2
MAPK: Mitogen-Activated Protein Kinase
MAPK: Mitagoen-activated kinase
MARK2: Serine/threonine-protein kinase MARK2
MEK: Mitogen-activated protein kinase kinase kinase
p-p38MAPK: phosphorylated p38MAPK
p38MAPK: p38 mitogen-activated protein kinase
PAK2: Serine/threonine-protein kinase PAK 2
pERK: phosphorylated ERK
PG: Plakoglobin
PKC: Protein Kinase C
PKP: Plakophilin
PKP2: Plakophilin 2
PKP3: Plakophilin 3
PLEC: Plectin
Raf: Rapidly Accelerated Fibrosarcoma
Ras: Rat sarcoma
RFP: Red fluorescent protein
RHOG: Rho-related GTP-binding protein RhoG
ROCK1: Rho-associated protein kinase 1
RUD: Repeat unit domain
Src: Proto-oncogene tyrosine-protein kinase Src
STK24: Serine/threonine-protein kinase 24
STRING: Search Tool for Retrieval of Interacting Genes/proteins
TJP1: Tight junction protein 1
TJP3: Tight junction protein 3
VEGFR: Vascular Endothelial Growth Factor
WB: Western blot
YAP1: Transcriptional coactivator YAP1

